# Development of an amplicon nanopore sequencing strategy for detection of mutations conferring intermediate resistance to vancomycin in *Staphylococcus aureus* strains

**DOI:** 10.1101/2022.07.31.502206

**Authors:** Abraham G. Moller, Robert A. Petit, Michelle H. Davis, Timothy D. Read

**Affiliations:** Division of Infectious Diseases, Department of Medicine, Emory University, Atlanta, GA, 30322; Microbiology and Molecular Genetics (MMG), Graduate Division of Biological and Biomedical Sciences (GDBBS), Emory University, Atlanta, GA, 30322; Theiagen Genomics, Highlands Ranch, CO, 80129

## Abstract

*Staphylococcus aureus* is a major cause of bacteremia and other hospital-acquired infections. The cell-wall active antibiotic vancomycin is commonly used to treat both methicillin-resistant (MRSA) and sensitive (MSSA) infections, but vancomycin intermediate *S. aureus* (VISA) variants can arise through *de novo* mutations. Here we performed pilot experiments to develop a combined PCR/long-read sequencing-based method for detection of previously known VISA-causing mutations. We amplified 16 genes (*walR*, *walK*, *rpoB*, *graR*, *graS, vraF*, *vraG*, *stpI*, *vraR*, *vraS*, *agrA*, *sarA*, *clpP*, *ccpA*, *prsA*, and *yvqF*) known from prior studies to be associated with mutations responsible for VISA as 10 amplicons and sequenced amplicon pools as long-reads with Oxford Nanopore adapter ligation on Flongle flow cells. We then detected mutations by mapping reads against a parental consensus or known reference sequence and comparing called variants against a database of known VISA mutations from laboratory selection. There was high (>1000x) coverage of each amplicon in the pool, no relationship between amplicon length and coverage, and the ability to detect the causative mutation (*walK* 646C>G) in a VISA mutant derived from the USA300 strain (N384-3 from parental strain N384). Mixing mutant (N384-3) and parental (N384) DNA at various ratios from 0 to 1 mutant suggested a mutation detection threshold of roughly the average minor allele frequency of 6.5% at 95% confidence (two standard errors above mean mutation frequency). The study lays the groundwork for direct *S. aureus* antibiotic phenotype inference using rapid nanopore sequencing from clinical samples.

**Importance:** Bacteremia mortality is known to increase rapidly with time after infection, making rapid diagnostics and treatment necessary. Successful treatment depends on correct administration of antibiotics based on knowledge of strain antibiotic susceptibility. *Staphylococcus aureus* is a major causative agent of bacteremia and is also increasingly antibiotic resistant. In this work, we develop a method to improve detection of a complex, polygenic antibiotic resistance phenotype in *S. aureus*, vancomycin-intermediate resistance (VISA) through long-read genomic sequencing of amplicons representing genes most commonly mutated in VISA selection. This method both speeds up VISA determination relative to purely culture-based detection and incorporates the most comprehensive database of VISA genetic determinants known to date.

## Introduction

*Staphylococcus aureus* is a commensal carried by 30% of humans (1) and a nosocomial pathogen that causes diverse pathologies. Many clinical strains are antibiotic resistant, with methicillin resistance (MRSA) first reported in the 1960s (2). The glycopeptide vancomycin, first released in 1958, has long been used for the treatment of severe MRSA infections (3). However, resistance to vancomycin has emerged in *S. aureus* through two distinct genetic processes. High level vancomycin resistance (VRSA), acquired through horizontal acquisition of the *vanA* gene, though detected in *S. aureus* in 2002 (4, 5), fortunately remains rare. Alternatively, vancomycin-intermediate resistance (VISA), the focus of this study, arises through *de novo* chromosomal mutations under antibiotic selection that are more often encountered in the clinic (3).

VISA is defined by a vancomycin MIC between 4 and 8 μg/mL, while heterogeneous VISA (hVISA) strains have a vancomycin MIC between 2 and 4 μg/mL (VRSA is > 16 μg/mL). Mutations associated with VISA are found in multiple sites in several key genes (6). The VISA phenotype includes a thickened cell wall, reduced autolysis, increased capsule, and increased D-alanylation of teichoic acids (3). The thickened VISA cell wall both reduces vancomycin diffusion and contains more free D-alanyl-D-alanine that can bind vancomycin, leading to reduction of vancomycin reaching the site of cell wall synthesis (the septum), and thus reduced sensitivity to the antibiotic (3). VISA strains furthermore have been shown to have reduced vancomycin susceptibility *in vivo* (3, 7) and to be associated with persistent bacteremia in clinical studies (3, 8–10). Despite its prevalence and significance, current VISA detection methods remain laborious and time-consuming (3, 11–13). VISA detection typically requires broth microdilution determination of minimum inhibitory concentration (MIC), while hVISA detection requires the more complex population analysis profile-area under the curve (PAP-AUC) assay (3). MIC and Etest assays require at least 24 hours and the PAP-AUC assay requires 48 hours incubation before resistance determination (3). Improving VISA treatment will thus require more rapid detection methods reducing or eliminating the time taken culturing.

Nanopore-based sequencing technologies offer a quicker alternative to culture-based tests for antibiotic resistance. DNA can be sequenced by pulling it through a protein nanopore, measuring the corresponding changes in current, and converting this current trace into a DNA sequence (14). While it has a higher per-base error rate (~10%) than short-read sequencing (e.g., Illumina), nanopore sequencing not only results in longer read length relative to other sequencing technologies (over 1 kb per read) but is also faster - sequencing data can be collected in real time, as early as the beginning of the sequencing run. Improvements in base calling have also led to reduced error rates, addressing its major weakness (15).

Recent studies have demonstrated that nanopore sequencing can detect antibiotic resistance genes in metagenomic samples (16–19) and distinguish antibiotic-sensitive from resistant strains far faster than culture-based methods, which typically require overnight growth (20, 21). A method for distinguishing carbapenem-resistant from sensitive *Klebsiella pneumoniae* through 16S rRNA present in culture under imipenem treatment only requires four hours of culturing (20). Another method that extrapolates antibiotic resistance from pneumococcal sequence type takes only five minutes (21). No nanopore method has yet been developed for direct detection of resistance-causing mutations from PCR amplicons, but such mutations have been detected in metagenomic sequence data collected from urine containing *Neisseria gonorrhoeae* (22). Additionally, genomic prediction of antibiotic resistance must still be calibrated thoroughly against culture-based methods (23).

Here we present a pilot approach to rapidly determine likely vancomycin-intermediate *Staphylococcus aureus* (VISA)-conferring mutations in *S. aureus* strains or clinical samples by coupling PCR and nanopore sequencing. This method offers a potential improvement on the current state of the art (e.g., qPCR for resistance genes or mutations, culture-based testing) because there are many possible mutations in many genes that may cause VISA, including mutations we have never seen before, which makes sequencing followed by bioinformatic mutation detection and interpretation necessary. To our knowledge, this is the first study aimed at detecting mutations (SNPs and small indels) linked to antibiotic resistance rather than whole genes through nanopore sequencing. We developed a PCR scheme to amplify 10 regions containing the 16 genes (*walR, walK, rpoB, graR, graS, vraF, vraG, stpI, vraR, vraS, agrA, sarA, clpP, ccpA, prsA*, and *yvqF*) most likely to contain VISA-conferring mutations based on previous work (6). We then developed a method to detect such mutations without culturing through sequencing, alignment against a reference, and comparison to a database of VISA-conferring mutations, taking far less than the 24+ hours necessary for culture-based detection. As a proof of principle, we have sequenced VISA amplicons from a parent strain (N384) and one mutant (N384-3) to detect VISA-conferring mutations. We also detected the mutation when its abundance was as low as 10% relative to the parent and detected VISA mutations directly from clinical VISA strains. Future work will attempt to amplify all regions together through a multiplex PCR reaction and detect VISA mutations in the shortest possible time through machine learning methods.

## Results

### Development of a nanopore sequencing VISA detection pipeline

We developed a strategy combining DNA extraction from a *S. aureus* isolate, PCR amplification of *S. aureus* genes often implicated in VISA, and nanopore sequencing of these amplicons to identify mutations. Our proposed pipeline is outlined in Figure 1. *S. aureus* may be isolated either directly (isolation from culturing) or indirectly (metagenomic DNA extraction) from a clinical sample such as a blood bottle. *S. aureus* populations cannot be assumed to be clonal; instead, they may include a mixture of the VSSA parental strain and at least one VISA mutant, if not more. Challenges must be addressed in five general areas - 1) sample isolation, 2) PCR of multiple unlinked genomic regions, 3) sequencing, 4) mutation calling and calling of VISA/VSSA based on mutation patterns, and 5) bioinformatic determination of whether potential VISA mutations had been previously characterized. Sample isolation must both produce enough DNA for PCR and remove any contaminating non-bacterial DNA that would lead to subsequent nonspecific PCR. The PCR method must amplify each fragment to similar relative abeyance with the lowest DNA input. Mutation calling must distinguish sequencing errors from VISA-causing mutations and allow for detection of subpopulations of unfixed VISA mutations. False positive errors would suggest spurious emergent resistance in clinical samples; false negative errors would miss a true resistant subpopulation.

**Figure 1:**
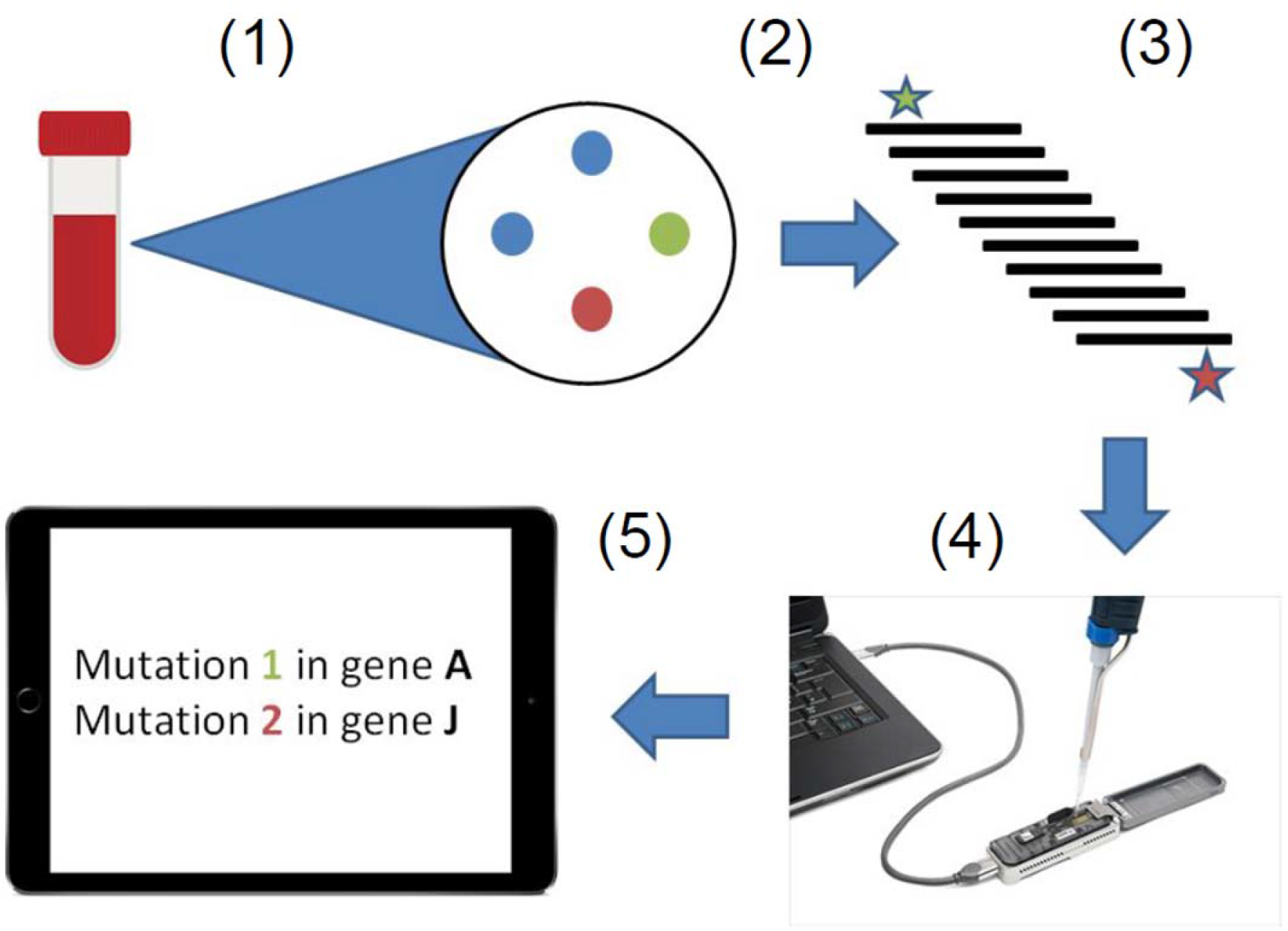
Overview of the VISA amplicon sequencing process, from sample isolation to SNP calling and comparison to known VISA mutations. Steps 1-5 represent sample isolation, DNA extraction, PCR, nanopore sequencing, and variant calling, respectively. Two VISA mutations are illustrated in red and green (steps 2, 3, and 5) while their parental strain is in blue.

### Steps 1 and 2 - sample isolation and PCR

Initial experiments only yielded appropriate DNA template concentrations for PCR from bacterial cultures themselves. We attempted spiking CD1 mouse blood with 1e2 or 1e4 CFU/mL N384 or N384-3, but in either case upon DNA extraction with the Qiagen DNeasy Blood and Tissue Mini Kit, we only obtained nonspecific amplification with PCR for all amplicons, presumably from mouse DNA (data not shown). We also found that microwaving culture pellets resulted in DNA too fragmented to amplify our long regions of interest. However, when we extracted DNA from isolated bacterial cultures, we did manage to amplify our regions of interest (Figure 2). We decided to forego development of blood-based extraction at this point to concentrate on Steps 2-4. We used DNA extracted from bacterial cultures with the modified Qiagen DNeasy Blood and Tissue Mini Kit protocol to serve as a PCR template in subsequent experiments (extracting DNA from mixed N384 and N384-3 cultures). With these templates (diluted to 0.2 ng/μL), we were able to amplify all 10 PCR amplicons (lengths in Table 2) to similar relative concentrations (70 ng/μL).

**Figure 2:**
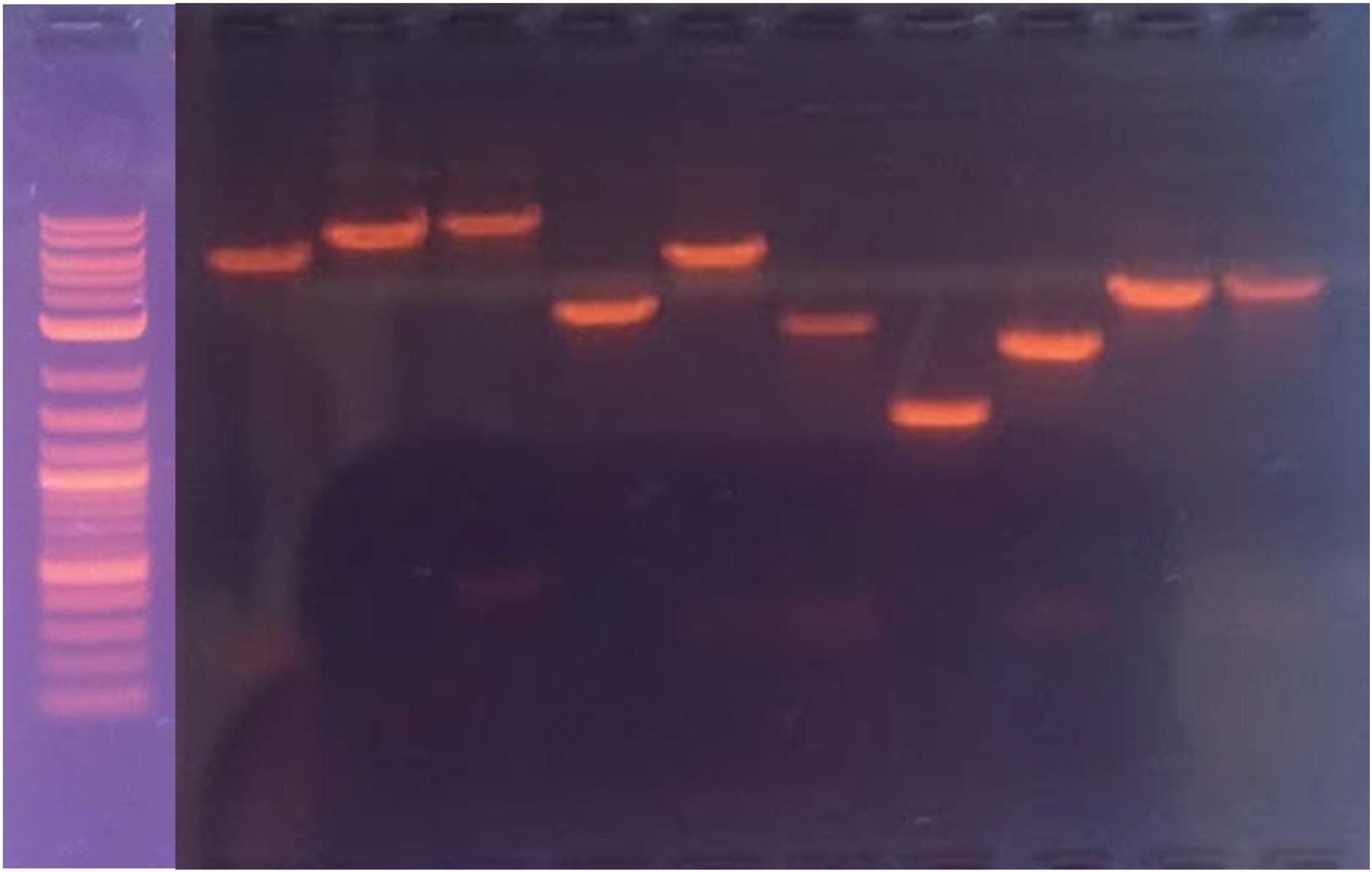
Evidence that our 10 VISA regions can be amplified from N384 mutant DNA extracted from pure culture. From left to right, the amplicons are *walRK, rpoB, graRS, vraFG, stpI, vraRS, agrA, sarA, clpP, ccpA, prsA*, and *yvqF*. The NEB 1 kb Plus ladder is shown for reference at left. The ladder and all amplicons were run at 110V for 30 minutes.

**Table 1:** Strains used in this study

| Strain | Description | Reference(s) |
| --- | --- | --- |
| N384 (NRS384) | Community-associated USA300 MRSA strain (CC8) | (33) |
| N384-3 | Laboratory selected VISA mutant in N384 background; mutation *walK* 646C>G Q216E | (33) |
| EUH15 | VSSA clinical isolate (CC5); vancomycin MIC is 1 µg/mL | (6) |
| 107 | VISA clinical isolate (CC5); vancomycin MIC is 4 µg/mL | (6) |

**Table 2:**
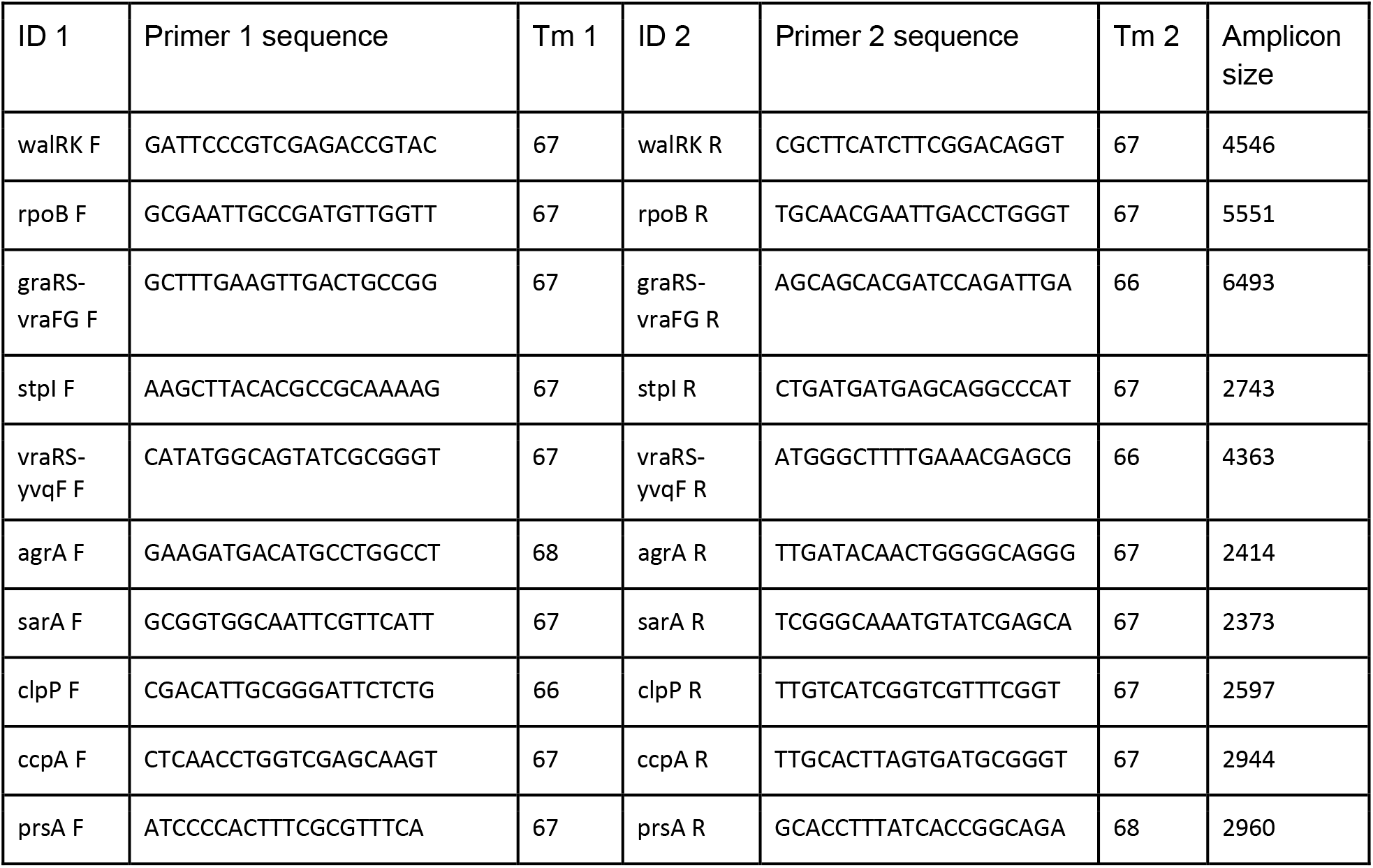
Primers used for amplifying each of 10 regions containing genes likely to contain VISA-conferring mutations, melting temperatures calculated for each primer with the NEB Tm calculator, and amplified region sizes.

### Steps 3 and 4 - sequencing and mutation detection: limit of detection approaches average minor allele frequency

To evaluate the limit of detection of our assay, we prepared three different mixes of parental VSSA (N384) and VISA (N384-3) mutant DNA - 1) culture mixtures, 2) DNA amplicon mixtures, and 3) simulated amplicon sequencing read mixes. We either mixed N384 and N384-3 liquid cultures, DNA amplicon pools of identical concentrations, or nanopore reads simulated from each genome in ratios from 100% parent to 100% mutant in 10% increments based on CFU/mL, DNA concentration (ng/μL), or ratio of simulated reads, respectively. We then called mutations through two different methods (bwa alignment (24) followed by bcftools consensus calling (25); medaka (26) alignment, consensus determination, and consensus variant calling) and analyzed aligned nucleotide counts for each mixture. Culture (cell) and DNA amplicon mixtures had similar coverages to each other but not to simulated DNA read mixtures based on non-parametric Wilcoxon tests (Figure 3A and 3B; p=5.5e-14 and 1.6e-10, relative to culture and DNA mixtures, respectively, in Figure 3B). The same pattern was present at a per-amplicon level (Figure 3A), except for *walRK*, where all three sets overlapped, and *prsA*, where the simulated coverage was significantly lower than that of either (p=6.3e-05 and 4.6e-4, relative to culture and DNA mixtures, respectively). In all cases (cell, DNA, or simulated mixtures), coverage was in excess of 3000-fold, which is well above the inverse of the error rate (~1/0.05 or 20-fold), suggesting coverage is high enough to compensate for at least random errors. Additionally, in all cases, there was no significant relationship between amplicon length and amplicon coverage (Figure 3C; p=0.94 for cell, p=0.087 for DNA, and p=0.19 for simulated mixtures, respectively).

**Figure 3:**
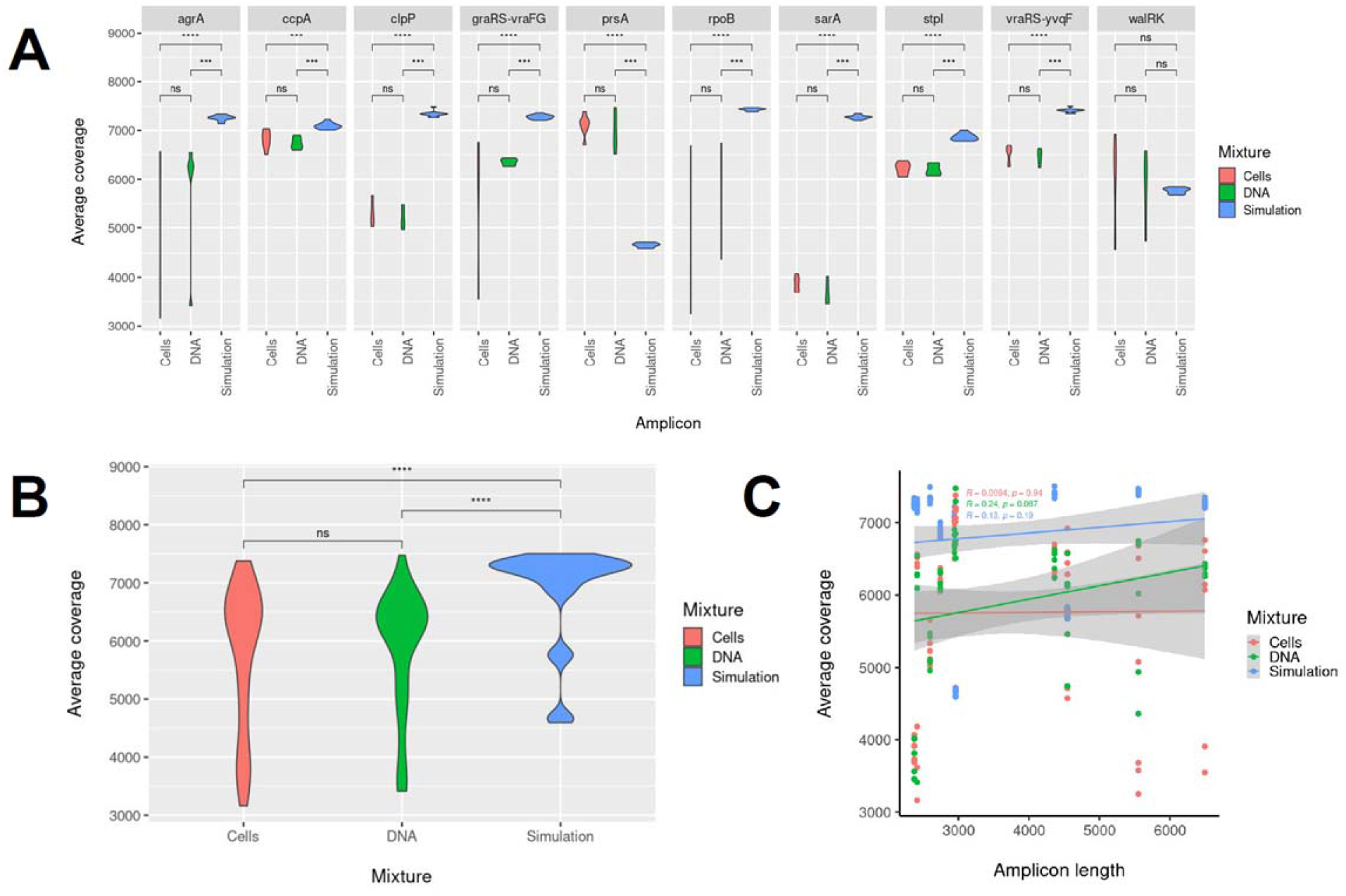
Gene coverage analyses for simulation (blue), DNA (amplicon; green), and cell (culture; red) parent/mutant mixtures. A) Coverage for each of the ten amplicons visualized as a violin plot; coverage for each amplicon is compared amongst mixture sets with a non-parametric WIlcoxon test. B) Coverage for each mixture set (simulation, DNA, or cell) visualized as a violin plot; coverage is compared between mixture sets with a non-parametric Wilcoxon test. C) Coverage against amplicon length for each mixture set presented with correlation and p-value.

Given that we achieved high coverage for all amplicons in all cases, we proceeded to call mutations in each mixture (Table 3) and analyze detected mutation frequency relation with mutant proportion in the mixture (Figure 4). Amongst the simulated mixtures, medaka only detected the N384-3 *walK* 646C>G mutation at 60% mutant proportion in the mutant/parent mixture or higher, while bwa/bcftools detected the mutation at a proportion of 50% mutant or higher (Table 3). For the cell and DNA mixtures, on the other hand, medaka detected this mutation at 50% mutant proportion or higher while bwa/bcftools detected it at 10% mutant or higher (Table 3). This suggests the standard variant calling pipeline (bwa/bcftools) was more sensitive with respect to mutant abundance than the faster medaka. In addition, bcftools had consistently lower coverage thresholds for detecting the *walK* mutation than medaka when subsampling read datasets (Table 5). Regarding the correlation between introduced and detected mutant proportion, we found the greatest deviation from a perfect correlation with the culture mixtures (Figure 4A). At 50% mutant, the detected mutant proportion was below that detected from all other mixtures at the same introduced mutant proportion. We hypothesize that differences in DNA extraction efficiency between the VISA mutant and VSSA strain explain this deviation, as the thicker VISA cell walls would make DNA harder to extract, leading to a lower detected mutation proportion than expected based on how the cultures were mixed.

**Figure 4:**
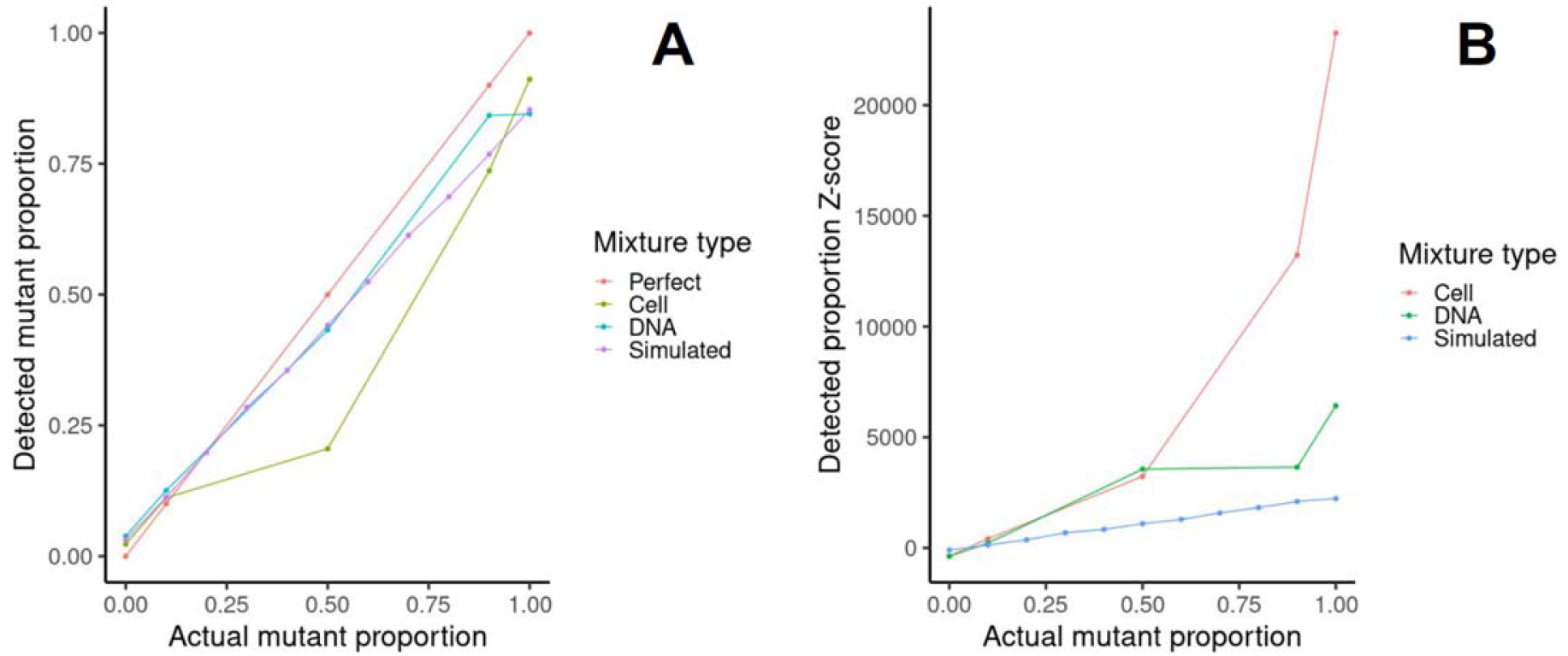
Evaluating limit of detection over a range of mutant/parent mixes (cultures, amplicon DNA, and simulated sequence reads). A) Introduced mutant proportion (x-axis) vs. detected mutation proportion (y-axis) for culture (olive-green), amplicon DNA (blue green), or simulated (purple) sequence read mixtures along with the diagonal expected if the introduced mutant proportion matched detected mutation proportion (perfect; red). B) Introduced mutant proportion (x-axis) vs. detected mutation Z-score (y-axis) for cell (red), amplicon DNA (green), or simulated (blue) sequence read mixtures.

**Table 3:** Mutation calls (whether *walK* 646C>G was called) by bwa/bcftools and medaka for simulated, amplicon DNA, and culture mixtures

| Proportion<br>N384/N384-3 | Simulated (NanoSim) | Amplicon DNA | Culture |
| --- | --- | --- | --- |
| 100% N384 | no/no | no/no | no/no |
| 90% N384<br>10% N384-3 | no/no | yes/no | yes/no |
| 80% N384<br>20% N384-3 | no/no |  |  |
| 70% N384<br>30% N384-3 | no/no |  |  |
| 60% N384<br>40% N384-3 | no/no |  |  |
| 50% N384<br>50% N384-3 | yes/no | yes/yes | yes/no |
| 40% N384<br>60% N384-3 | yes/yes |  |  |
| 30% N384<br>70% N384-3 | yes/yes |  |  |
| 20% N384<br>80% N384-3 | yes/yes |  |  |
| 10% N384<br>90% N384-3 | yes/yes | yes/yes | yes/yes |
| 100% N384-3 | yes/yes | yes/yes | yes/yes |

**Table 4:** Mutation calls by bwa/bcftools and medaka for test strains EUH15 and 107. Mutations and reference coordinates are given relative to USA300 FPR3757.

| Strain | bwa/bcftools | medaka |
| --- | --- | --- |
| 107 (VISA) | WalK A243T (26392)<br>GraR E15K (719060)<br>VraG T217I (718728)<br>VraS A314V (2026309) | WalK A243T (26392)<br>GraR E15K (719060)<br>VraS A314V (2026309) |
| EUH15 (VSSA) | none | none |

**Table 5:**
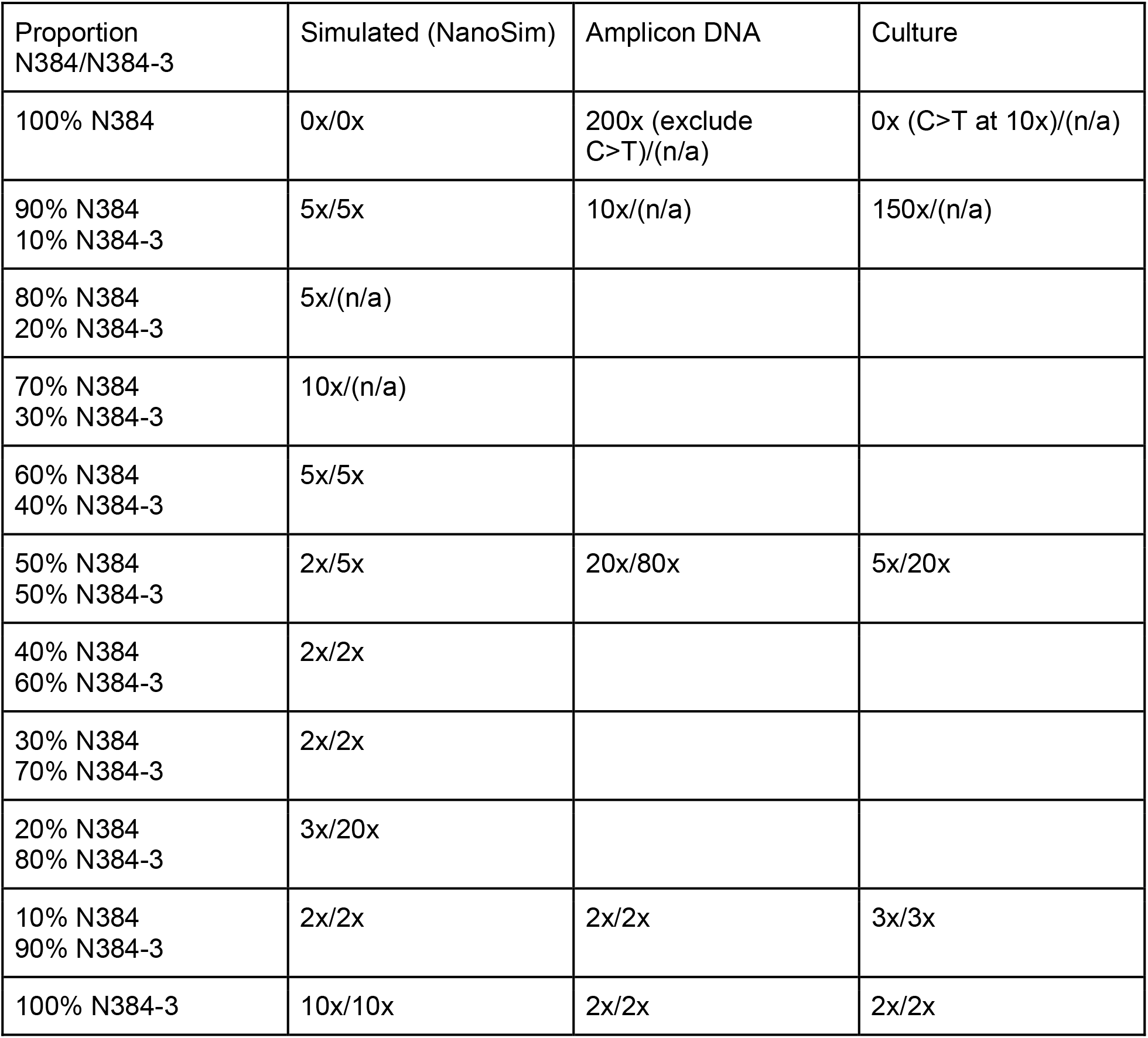
Minimum coverage required to call *walK* 646C>G mutation with bwa/bcftools and medaka (listed first and second in each column) in mutants or mutant mixtures or minimum coverage required to exclude mutation from wild-type samples for simulated, amplicon DNA, and culture mixtures. Cases in which the mutation was never detected are listed as n/a.

**Table 6:** Minimum coverage required to call *walK, graR, vraG*, and *vraS* mutations with bwa/bcftools and medaka in VISA clinical strain or minimum coverage required to exclude mutations from wild-type samples for test strains EUH15 and 107. Mutation calls by bwa/bcftools and medaka given for test strains EUH15 and 107. Mutations and reference coordinates are given relative to USA300 FPR3757.

| Strain | bwa/bcftools | medaka |
| --- | --- | --- |
| 107 (VISA) | WalK A243T (26392) (2x)<br>GraR E15K (719060) (2x)<br>VraG T217I (718728) (>500x)<br>VraS A314V (2026309) (2x) | WalK A243T (26392) (2x)<br>GraR E15K (719060) (2x)<br>VraS A314V (2026309) (5x) |
| EUH15 (VSSA) | None<br>WalK (26392) (20x)<br>GraR (719060) (50x)<br>VraG (718728) (0x)<br>VraS (2026309) (80x) | None<br>WalK (26392) (0x)<br>GraR (719060) (0x)<br>VraG (718728) (0x)<br>VraS (2026309) (0x) |

We also examined the Z-score for each mixture and each set to determine the limit of detection based on average and standard error of inherent sequencing variation found in the alignments (Figure 4B). In all cases, average minor (non-consensus) allele frequency was roughly 6.5% and tested mutant proportions above this threshold were well more than two standard errors (in fact, at least 100 standard errors) above the mean. Z-scores were highest for cell mixtures amongst the sets of mixtures evaluated (DNA and simulated reads). This suggests that the limit of detection based on our method is close (standard errors of minor allele frequency were roughly 1e-4) to the average minor allele frequency.

### Successful identification of VISA-associated mutations in a clinical VISA strain

To show our assay could identify VISA mutations in clinical strains, we sequenced two known clinical strains - one VSSA (EUH15) and one VISA (107) - characterized in a previous study. Strain 107 is known to contain four missense mutations in VISA-associated genes (WalK A243T at 26374, GraR E15K at 708287, VraG T217I at 711484, VraS A314V at 1947464) relative to the N315 reference, while strain EUH15 contains none of these four. Mutation calling results from two different methods (bwa/bcftools and medaka) are shown in Table 4. The first method (bwa/bcftools) identified all four mutations in strain 107 but not strain EUH15, but the second method (medaka) identified only three (in *walK, graR*, and *vraS*). This indicates that we can distinguish VISA from VSSA in at least two unknown clinical strains with our assay and bioinformatic pipeline. Standard variant calling (bwa/bcftools) outperformed rapid variant calling (medaka) in coverage sensitivity as we observed in our limit of detection study. As noted for the N384 parent/mutant mixtures, both medaka and bcftools detected VISA mutations of interest at 5x coverage or less, except for the VraG T217I mutation (500x for bcftools).

### Step 5 - bioinformatic analysis: construction of a VISA-associated mutation database to enhance detection

In order to broaden the scope of our method, we built a curated database of all VISA-associated mutations to make clinical VISA detection as sensitive and specific as possible. We collected mutations in the sixteen genes of interest with four levels of support for VISA causation - 1) laboratory selection for VISA, 2) identification in clinical VISA strains but not VSSA strains, 3) variation in our curated Staphopia database (27) that shows the potential for VISA causation (e.g., nonsynonymous mutations), and 4) simulated mutations in our VISA-associated genes with potential for VISA causation. We noted the most mutations in Staphopia and synthetic mutation classes in the longest genes, such as 3551 bp *rpoB* (10653 synthetic mutations), 1890 bp *vraG* (5670 synthetic mutations), and 1826 bp *walK* (5478 synthetic mutations) (Figure 5A). However, in the lab and clinical classes, *vraG* instead had the most mutations identified (229 lab and 217 clinical, respectively), followed by *graS* (112 and 108) and *vraF* (78 and 80). We also found no mutations in *sarA* called in either lab or clinical datasets (Figures 5A and 5B). We then annotated mutation calls with SnpEff to examine the distribution of mutation types (low, moderate, and high effect, corresponding to mainly synonymous, missense, and nonsense mutations, respectively) in each class. We found the most consistent mutation type distribution in the synthetic mutation class (20.50±0.94% low, 73.43±0.82% moderate, and 6.07±0.72% high; mean and standard deviation across genes, respectively). Low effect mutations were far more common in every other mutation class (47.76±10.32% in the Staphopia class, 58.48±27.24% in the lab class, and 70.86±23.3% in the clinical class), by statistically significant margins in all cases (Paired t-tests; p = 1.103e-8 for Staphopia, 4.807e-5 for lab, and 2.81e-7 for clinical classes).

**Figure 5:**
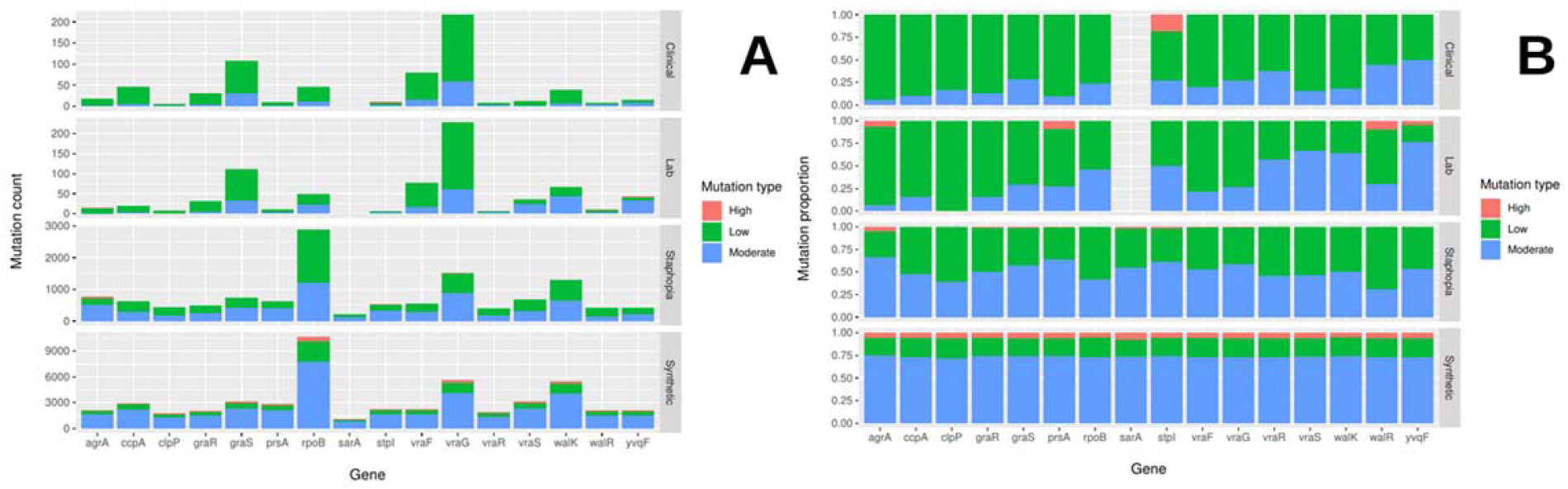
Distributions of annotated variant types in VISA mutation database. Variants in 16 known VISA-associated genes were called in clinical VISA and lab-adapted VISA short read data against the Staph_ASR reference with snippy and annotated with SnpEff. Variants in our curated database of 40,000+ *S. aureus* genomes and all possible simulated mutations in the VISA-associated genes were also annotated with SnpEff. Proportions of high, moderate, and low effect mutations are marked in red, blue and green, respectively. Either raw mutation counts (A) or proportions (B) are shown for each gene and mutation effect class, respectively.

The critical control to evaluate our ability to discriminate VISA from VSSA based on this database was the overlap between these VISA database mutations and mutations called from parental N384 nanopore reads against the N384 reference genome. Such mutations would likely represent systematic sequencing errors that would be called false positives. To determine such overlap, we compared N384 and N384-3 sample mutation calls against the Staph_ASR reference (27) to each other and N384 or N384-3 unique calls to each of the four VISA mutation databases, reporting results in Table 7. We found that the number of shared mutations depends on the mutation detection method (bcftools or medaka). Neither detection method identified any mutations in the N384 parent that were also found in clinical or lab strains, but bcftools did call 44 mutations shared with those found in Staphopia strains (9 low, 33 moderate, and 2 high). In N384-3, on the other hand, medaka only detected the expected missense SNP in *walK* (646C>G, Q216E) and a conservative in-frame deletion in *vraR* (418T>TTTTTCATACGGTTACGCA, duplication of AAs 134-139), both of which were also found in the lab-adapted strain mutation database. Meanwhile, bcftools called over 100 mutations shared with clinical, lab, and staphopia mutations. In each case, over 100 of these mutations were moderate effect (126 shared clinical, 131 shared lab, and 103 shared Staphopia mutations, respectively). This suggested that medaka is more specific while bcftools is more sensitive in its mutation calling.

**Table 7:** Mutations bwa/bcftools or medaka (first and second counts in each cell for each mutation type, respectively) called against Staph_ASR shared between N384 or N384-3 and VISA mutation databases (clinical, lab, and Staphopia) segregated by annotation type (low, moderate, or high effect) for each shared group. Mutations were annotated with SnpEff.

| Database | N384 | N384-3 |
| --- | --- | --- |
| Clinical | 0/0 | 7/0 low<br>126/0 moderate<br>6/0 high |
| Lab | 0/0 | 8/0 low<br>131/2 moderate<br>5/0 high |
| Staphopia | 9/0 low<br>33/0 moderate<br>2/0 high | 41/0 low<br>103/0 moderate<br>1/0 high |

## Discussion

Nanopore sequencing may revolutionize sequence-based antimicrobial resistance detection as it allows very rapid data generation. Multiple studies have shown that nanopore sequencing can rapidly (15 minutes or less after sequencing begins) identify antibiotic resistance genes in plasmids (28) or antibiotic resistant strain lineages (21) and detect antibiotic resistance mutations in metagenomic sequencing from clinical samples (22). In this study, we explored PCR amplification of a set of genes known to be commonly associated with vancomycin-intermediate resistance. We took advantage of the long-read capabilities of the technology to reduce the number of amplicons to only 10 and showed that we could rapidly acquire 3000x coverage from each. This approach offers the potential advantage of not needing to culture *S. aureus* from clinical samples, which could take 24-48 hours. Indeed, the time lost to culturing has been suggested to be a cause of mortality itself, as mortality increases 10% with each hour in septic shock cases (29). Another advantage of the PCR approach is that sequence data returned is only the portions of the pathogen genome known from prior studies to be associated with the phenotype, rather than metagenomic, or even whole genome data.

These results suggest that the assay has potential to identify VISA mutations in unknown clinical strains but work still must be done to further improve 1) DNA extraction, 2) PCR amplification, and 3) basecaling.

In regard to the first area, we were unable to extract sufficiently high quality DNA for downstream steps. Attempts to amplify VISA amplicons directly from mouse blood spiked with 1e2 or 1e4 CFU/mL *S. aureus* failed with extensive nonspecific amplification. This suggests at least one culturing step may be necessary to isolate *S. aureus* for DNA extraction and PCR. We must reduce culturing time to as little as necessary to get enough isolated bacterial culture, followed by quick enough (~1 hr, including 30 min lysostaphin/lysozyme treatment) DNA extraction. The development of rapid approaches to extract bacterial DNA from clinical samples should be considered a critical research area. Recent advances have reduced human DNA 10-fold while maintaining bacterial DNA in a 7 hour biopsy to pathogen identification protocol (30) and identified DNA extraction kits (Qiagen UCP pathogen mini kit) that best depleted human DNA relative to *Neisseria gonorrhoeae* DNA in clinical urine samples (31).

The PCR primers amplified all 10 regions in simplex reactions that took about 2 hours, but further development would be necessary for optimization. Ideally, these reactions should be multiplexed in as few tubes as possible. The solution may be to explore using a higher number of shorter amplicons. This scheme may also be more robust to lower *S. aureus* DNA template yield and quality. The limits of detection of the PCR system need to be thoroughly explored on spiked and real clinical samples. At the very least, the PCR and sequencing should be able to verify the presence of *S. aureus* in the clinical sample to a certain CFU/mL titer.

Regarding sequencing time, we need to determine the minimum number of reads necessary to call a mutation in our mutant/parent mixtures by randomly subsampling various numbers of reads and repeating our variant calling pipelines. We also need to see whether fast basecalling is sufficiently accurate to call VISA mutations, as we used the high accuracy, neural network-dependent bonito basecaller (32) with external GPUs after sequencing. We attempted to speed up alignment with medaka, but our standard protocol (bwa/bcftools) sensitively and completely called VISA mutations. Nanopore technology currently has higher sequencing error (~3-5% average per read per base for the newest versions) than other methods. In addition, this sequencing error is often systematic, focused in loci such as homopolymer regions. The critical problem to overcome for these nanopore applications is distinguishing sequencing errors from causative resistance mutations when analyzing sequence traces. Medaka, a rapid neural network-based alternative to standard alignment and variant calling (26), proved ineffective both in sensitively detecting mutations in the mutant/parent mixtures and in comprehensive detection of the strain 107 VISA mutations.

Future work will involve validating this assay further in more genetically diverse VISA and VSSA strains and further correcting for nanopore sequencing error. The construction of a VISA mutation database for comprehensive mutation detection in unknown samples, including lab-evolved, clinical, and predicted VISA mutations in one database, has already provided insights into the mutation call sensitivity and specificity of our assay. We anticipate this database should make detection possible across diverse strains and serve as a resource for VISA resistance characterization and population genetics more generally. The critical control we must resolve is sequencing errors that overlap with mutations in this database. We have assessed this by comparing N384 parental mutations with those in our database. We expect that further future improvements in nanopore basecalling and mutation calling (with medaka) will translate into reductions in false positive VISA calls in our assay.

## Materials and Methods

### Strain selection and media

Either a MRSA parent strain (N384) and its lab-selected VISA mutant (N384-3) (33) or clinical VSSA (EUH15) or VISA (107) strains were used for all downstream experiments. N384-3 was selected from N384 through stepwise evolution in vancomycin up to 8 μg/mL. Strains and associated metadata (vancomycin MICs) are listed in Table 1. All strains were grown on tryptic soy agar (TSA) or in tryptic soy broth (TSB) at 37°C with 225 rpm agitation for liquid culture.

### Preparation of mutant/parent mixtures (cell and DNA) for evaluating limit of detection

In order to evaluate the limit of mutation detection, mutant and parent sequence were mixed together in five ratios from 100% N384-3 mutant to 100% N384 parent (100% N384-3, 90% N384-3; 10% N384, 50% N384-3; 50%N384, 10% N384-3; 90% N384, 100% N384). Either turbid bacterial cultures or amplicon DNA volume were mixed in these ratios (total amplicon volume of 100 μL; 1 mL culture volume; 3e9 CFU/mL). DNA (including amplicon) concentrations were measured with a Qubit dsDNA Broad Range (BR) Assay Kit (Invitrogen). DNA was extracted from cell mixtures with a modified Qiagen DNeasy Blood and Tissue Mini Kit. Cultures were pre-treated for 30 minutes with 0.2 mg/mL lysostaphin and 1 mg/mL lysozyme at 37°C to cleave *S. aureus* cell walls and then DNA was extracted following manufacturer’s directions.

### Amplifying VISA-associated loci

VISA amplicons represented regions from the USA300 FPR3757 genome containing VISA-associated genes or clusters of such genes (e.g., the *walRK* two-component system) and 1000 bp adjacent sequence on either side (Table 2). These regions were amplified through PCR from N384 parent/mutant DNA mixture templates (0.2 ng/μL). All regions were amplified using NEB Q5 High-Fidelity 2X Master Mix and the following PCR program: 98°C for 30 seconds (initial denaturation); 98°C for 10 seconds, 65°C for 30 seconds (annealing), and 72°C for 3 minutes 30 seconds (35 cycles; extension for up to 6.5 kbp sequence); 72°C for 2 minutes, final extension; and then hold at 4°C.

### Amplicon nanopore sequencing

VISA amplicons were pooled to a total of 1 μg for sequencing (equal quantities of each amplicon). VISA amplicon libraries were generated using the 1D ligation sequencing kit (SQK-LSK109; Oxford Nanopore) modified for amplicon sequencing. 0.2 pmol of amplicon DNA was ligated to adapters to bias ligation reactions toward individual amplicons. Libraries were sequenced on Flongle FLO-FLG001 flow cells (Oxford Nanopore) on a MinION instrument. Read data (event-level, FAST5 format) was collected on a PC using MinKNOW software (Oxford Nanopore). Raw sequencing data generated from the MinION were deposited in the NCBI Sequence Read Archive (SRA) under BioProject accession PRJNA863907.

### Basecalling, alignment, and variant calling

For *de novo* basecalling, bonito (version 0.3.6) (32) was used with nucleotide output stored as a FASTA file. For detecting nucleotide changes compared to a reference (mapping), all basecalled reads called by the bonito software (including EUH15 and 107 reads) were aligned against the USA300 FPR3757 reference using BWA (version 0.7.17; parameter -x ont2d) (24). Variants were called with bcftools (version 1.11; mpileup/call on the resulting read alignment with consensus and multiallelic variant calling) and medaka (version 1.6.0; default parameters, reads against USA300 FPR3757 reference), with results saved in a VCF file. Variants were further filtered down to biallelic sites with bcftools view. The consensus sequence was determined using samtools mpileup (-B option selected to cancel base alignment quality assessment) and bcftools (-c option for consensus determination). Variants were compared to the known N384-3 VISA mutations (*walK* 646C>G) in the case of N384 parent/mutant mixtures and known, previously identified VISA-associated mutations in the case of EUH15 and 107 test strains.

### *In silico* read simulation and mutation frequency analysis

*In silico* amplicon sequence mixtures (300,000 reads in each case; 100% N384-3, through 100% N384 in 10% increments of each) were simulated using NanoSim-H (version 1.1.0.4) (34, 35) using default parameters from the wild-type (N384) and VISA mutant (N384-3) amplicon sequences concatenated into a single FASTA sequence (predicted amplicons from primers in Table 2). Amplicon reads were simulated with an error profile trained from bonito-basecalled N384 parent amplicon (sequenced from amplicon DNA rather than cell mixture series). A total of 300,000 reads were simulated in each case. These read mixtures were then used together with cell and DNA mixture reads to determine how VISA mutation frequency correlated between variant calling (observed mutation frequency) and VISA proportion (predicted VISA mutation frequency) and determine the detection threshold without use of medaka. In order to address these questions, simulated and real (cell or DNA) mixtures were aligned against the USA300 FPR3757 (N384) reference using BWA (version 0.7.17; parameter -x ont2d) (24) and analyzed for variant counts with samtools (25) mpileup (-B option selected to cancel base alignment quality assessment). Mpileup output was further processed into a table with counts of every base, positive strand matches (.), negative strand matches (,), and overall mapped read coverage at each amplicon site. Overall non-reference nucleotide frequency was calculated for every mapped amplicon nucleotide to generate an overall distribution with a mean and standard deviation. For each proportion mutant, the detected mutation (*walK* 646C>G at USA300 FPR3757 position 26311) is proportional to mapped coverage and the Z-score. The detection limit was calculated as the extrapolated observed mutation proportion for which the Z-score exceeded two (two standard deviations above the mean to give two-tailed 95% confidence).

### Evaluating minimum coverage necessary for VISA mutation detection

We randomly subsampled three types of nanopore amplicon data (in silico simulated, cellular, and DNA parent/mutant mixtures) plus the additional test VSSA (EUH15) and VISA (107) strains to sequence coverages ranging from 2 to 500x in order to detect the minimum sequence coverage necessary for correctly detecting VISA mutations. We subsampled reads with seqtk (version 1.3-r106). Subsampled reads were aligned against the USA300 FPR3757 reference with bwa and mutations were called with bcftools and medaka as previously described. Subsampled read mutation calls were then compared to the expected mutation (*walK* 646C>G at USA300 FPR3757 position 26311). Minimum coverage necessary to call this mutation was estimated as the minimum coverage for which the expected mutation was detected in the mutation calls.

### Constructing a VISA mutation database

A database of VISA mutation was constructed from three levels of information: 1) lab-evolved VISA mutations, 2) VISA-associated mutations in clinical strains, 3) predicted VISA-causing mutations detected in our database of 40,000+ *S. aureus* genomes and 4) simulated mutations in VISA-associated genes expected to cause VISA. This database was constructed by 1) collecting mutations identified in our laboratory VISA evolution studies (33), 2) collecting those previously identified in clinical VISA strains in our lab (6), 3) finding nonsense or missense mutations relative to the N315 reference in our Staphopia database (27), and 4) performing a saturating mutagenesis of the genes in our amplicons and identifying all nonsense or missense mutations predicted to confer VISA. Previously screened clinical strains were selected as VISA if they showed an MIC over 4 μg/mL in any MIC determination method used (broth microdilution - BMD, Etest, or population analysis profile-area under the curve - PAP-AUC). Short reads for lab-adapted VISA strains, clinical VISA strains selected based on the stated MIC criteria, or all strains in the Staphopia database were aligned against the Staph_ASR reference with bwa. Mutations were called against the Staph_ASR reference (classes 1 and 2 described previously) with snippy (version 4.6.0) (36), selected from the Staphopia database (27), or simulated in the Staph_ASR reference (class 4) with the syntheticVCF tool compiled in C (37). Mutations called per strain in classes 1-3 were then merged into a nonredundant set per class with bcftools (version 1.11) merge. Variants were filtered for VISA gene coordinates with vcftools (version 0.1.16) and variant effects were annotated with SnpEff (version 5.0e). The Staph_ASR genome was annotated with bakta (version 1.4.0) (38) for use in SnpEff.

### Comparison of nanopore sample mutation calls against VISA mutation database

To evaluate sensitivity and specificity of mutation calling from our nanopore samples (N384 and N384-3), we aligned N384 and N384-3 (100% of each in the amplicon DNA series) nanopore reads against the Staph_ASR genome with bwa and called variants with bcftools call or medaka. We then identified medaka or bcftools-called mutations specific to N384 or N384-3 by comparing N384 and N384-3 mutation calls with bcftools isec. N384 or N384-3 specific mutations were then compared to lab, clinical, and Staphopia mutation databases also with bcftools isec. Mutations found to be shared between N384 or N384-3 specific mutations and each database were then annotated with SnpEff and classified by mutation type (low, moderate, or high effect).

## Acknowledgements

We thank members of the lab for constructive criticism. Timothy D. Read was supported by NIH R21 AI 138079-02, while Abraham G. Moller was supported by the NSF GRFP. We thank NSF for providing GPU resources via the XSEDE supercomputing consortium to conduct high accuracy basecalling with bonito.

